# Robust Trachea Segmentation from CT Imaging Using Fully Automated and Prompt-Based Models

**DOI:** 10.64898/2026.01.16.699540

**Authors:** E. Toulkeridou, Z. Antoniou, A. Panayides

## Abstract

Accurate trachea segmentation from computed tomography (CT) is a prerequisite for image-guided airway assessment, toward precision tracheostomy planning and safe endotracheal tube placement. Trachea-specific delineation remains challenging due to elongated geometry, small cross-sectional area, partial-volume effects, motion-trigerred artifacts, and heterogeneous surrounding tissues. Here, we study two complementary segmentation paradigms: a fully automated, self-configuring baseline (nnU-Net) and a prompt-conditioned foundation model (Med-SAM) derived from the Segment Anything Model (SAM). We evaluate both approaches on two heterogeneous regimes: (i) a spatially consistent volumetric dataset (AeroPath) and (ii) a large-scale, slice-based dataset without reliable volumetric continuity (OSIC). We further propose a hybrid inference strategy that enables fully automated prompting by deriving bounding-box prompts for MedSAM from coarse nnU-Net predictions. Results show that dataset structure critically influences reliability, interpretability, and deployment feasibility: volumetric continuity benefits fully automatic segmentation, while prompt-conditioned inference improves ROI-constrained interpretability in slice-based settings but introduces prompt sensitivity. We discuss limitations and outline clinically grounded evaluation and EHR integration directions for precision airway management.

## 1 Introduction

Airway management procedures, including tracheostomy and endotracheal tube (ETT) placement, are essential interventions in intensive care, emergency medicine, and surgical practice [1, 2, 3, 4, 5]. Despite their frequency, they remain technically demanding and can lead to severe complications when anatomical variability, pathology, or inaccurate placement occurs [6, 3]. Inadequate assessment of tracheal anatomy can result in airway trauma, tube malpositioning, impaired ventilation, and prolonged hospitalization [1, 2].

Computed tomography (CT) plays a central role in airway assessment by providing high-resolution visualization of the trachea and surrounding structures [7, 8]. However, transforming raw CT into clinically actionable geometry requires accurate image segmentation [9, 10]. Manual delineation is time-consuming and prone to inter-/intra-observer variability [11]. Automated trachea segmentation is therefore a key enabling technology for image-guided airway assessment and precision airway management [7, 8].

Trachea-specific segmentation presents challenges that differ from broader lung or airway tree segmentation tasks.Unlike broader lung or airway tree segmentation, the trachea is narrow, elongated, and occupies a small fraction of the field-of-view. It exhibits substantial variability between patients (shape, diameter, orientation) and is sensitive to partial-volume effects, motion artefacts, and acquisition variability [7, 9, 10]. These properties increase dependence on spatial context and anatomical priors, and make clinically relevant downstream descriptors (e.g., diameter profiles or cross-sectional area) sensitive to small boundary errors even when overlap metrics appear strong [10].

Deep learning has transformed medical image segmentation [11, 12]. Classical fully convolutional models (FCN, U-Net and variants) established the modern baseline [13, 14, 15, 16]. Volumetric networks (e.g., V-Net) improved 3D consistency [17]. More recently, nnU-Net emerged as a strong reference by automatically adapting preprocessing, architecture, and training to dataset properties [18]. In parallel, transformer-based and hybrid architectures (TransUNet, Swin-based models) further advanced segmentation performance [19, 20, 21, 22].

SAM introduced prompt-conditioned segmentation at scale [23]. MedSAM adapts this paradigm to medical imaging, enabling ROI-constrained segmentation via prompts such as bounding boxes [24]. Bounding boxes are particularly well suited for elongated anatomy because they provide a compact spatial prior that reduces sensitivity to local ambiguities compared to point prompts [23, 24]. However, prompt-based inference introduces an explicit dependency: segmentation quality can become sensitive to prompt quality and tightness [24].

Despite rapid progress, the relative behavior of prompt-based foundation models vs. fully automatic pipelines for anatomically constrained segmentation under heterogeneous dataset structure remains under-explored [25, 26, 27]. In particular, slice-based archives without reliable volumetric continuity may constrain context exploitation and can affect qualitative traceability of predictions (e.g., slice re-indexing in preprocessing pipelines) [18].

We present:

- A systematic comparison of fully automatic (nnU-Net) and prompt-based (MedSAM) paradigms for trachea segmentation [18, 23, 24].
- An analysis of how dataset structure (volumetric vs. slice-based) impacts performance, interpretability, and deployment constraints [9, 10, 26].
- A hybrid inference strategy enabling fully automatic prompt-based refinement: nnU-Net provides coarse localization and generates bounding-box prompts for MedSAM [18, 24].

## 2 Methodology

### 2.1 Datasets and data representation

#### AeroPath (volumetric CT)

AeroPath consists of 27 high-resolution 3D chest CT volumes (NIfTI) with voxel-wise airway annotations including the trachea. Volumes exhibit consistent spacing and alignment, enabling context-aware learning across 7680 slices (beneficial for elongated anatomy) [28, 17, 18].

#### OSIC (slice-based CT)

OSIC comprises axial 16708 CT slices paired with segmentation masks but lacks reliable volumetric continuity (incomplete stacks, inconsistent spacing, uncertain slice ordering). We treat OSIC strictly as 2D samples to avoid introducing artificial adjacency assumptions [29, 9, 10].

### 2.2 Preprocessing and sample selection

#### Intensity processing

Raw CT intensities were clipped to a clinically relevant Hounsfield Unit range to reduce scanner-specific artifacts and extreme outliers. Each slice was subsequently normalized to the [0, 1] range using min–max normalization, performed independently to account for acquisition variability [18, 30].

#### Foreground filtering

Slices with empty trachea masks are excluded during training and evaluation to reduce severe class imbalance and avoid trivial background learning [31, 10].

#### Spatial handling

nnU-Net uses its automated resampling and intensity handling pipeline [18]. MedSAM inputs are resized to 1024 *×* 1024; masks are resized with nearest-neighbor interpolation to preserve label topology [23, 24].

### 2.3 Fully automatic segmentation with nnU-Net

nnU-Net is a self-configuring segmentation framework that automatically adapts network architecture, preprocessing, and training schedules to dataset-specific properties [18]. For AeroPath, volumetric configurations were employed to leverage spatial context across slices. For OSIC, a strictly 2D configuration was used to respect the slice-based nature of the data. Training employed a composite Dice and cross-entropy loss, with class balancing handled internally by nnU-Net. Standard data augmentation techniques, including random rotations, scaling, and intensity perturbations, were applied using default configurations [18] [30].At inference time, nnU-Net produces segmentation masks in a fully automatic manner without requiring auxiliary inputs or user interaction. We optimize using Adam and standard deep learning training practice [32, 33].

### 2.4 Prompt-based segmentation with MedSAM (bounding boxes)

MedSAM is a medical adaptation of the Segment Anything Model, consisting of an image encoder, prompt encoder, and mask decoder. In this work, bounding box prompts were used exclusively [23, 24]. We use bounding boxes as prompts because they provide a robust spatial prior for small elongated targets [24]. During training, bounding boxes were derived from ground truth masks. We froze the image encoder and fine-tuned prompt encoder and mask decoder using Dice and BCE losses [24, 31].

### 2.5 Hybrid inference pipeline

To avoid manual prompting at inference time, we used hybrid prompting: (1) nnU-Net predicts a coarse trachea mask, (2) a bounding box is extracted, and (3) MedSAM refines the segmentation conditioned on the box [18, 24]. This combines automatic global localization with ROI-constrained refinement.

### 2.6 Training Setup and Hyperparameters

#### AeroPath

For the AeroPath dataset, segmentation was performed using both a volumetric convolutional architecture and a foundation-model–based approach, reflecting complementary design choices.

AeroPath CT scans were processed and segmented as full 3D volumes using the nnU-Net 3D full-resolution configuration. The network was trained on volumetric patches of size 128 × 128 × 128 voxels, automatically determined by nnU-Net’s self-configuring pipeline based on dataset statistics. Input images were resampled to a common target spacing derived from the median voxel spacing of the training set, and intensity normalization was applied using nonzero masks.

Training followed nnU-Net’s standard five-fold cross-validation protocol; however, folds were trained independently and stored separately to ensure robustness against interruptions. The network was optimized using stochastic gradient descent with Nesterov momentum, an initial learning rate of 0.01, and a polynomial learning rate decay schedule. Mixed-precision training was enabled to reduce memory usage and accelerate convergence. Each fold was trained for 30 epochs, with intermediate checkpoints saved to allow resumption in case of early termination.

In parallel, AeroPath scans were decomposed into axial 2D slices for training with MedSAM, as the model operates natively in 2D. The model was fine-tuned from pretrained MedSAM weights using slice-level supervision derived from the volumetric ground-truth annotations. Bounding box prompts tightly enclosing the trachea were used to guide segmentation. Training was performed for 30 epochs, with checkpoints saved at every epoch. Final 3D segmentations were obtained by stacking per-slice predictions along the axial axis.

#### OSIC

Due to the heterogeneous acquisition protocols and large inter-slice spacing present in the OSIC dataset, reliable volumetric reconstruction was not feasible. Consequently, all experiments on OSIC were conducted strictly in 2D, for both nnU-Net and MedSAM.

OSIC CT scans were processed as independent axial slices and used to train the nnU-Net 2D configuration. Slice-wise preprocessing, normalization, and augmentation were automatically configured by nnU-Net. Training employed stochastic gradient descent with momentum and nnU-Net’s default loss formulation (Dice + cross-entropy). Models were trained for 30 epochs, and validation was performed at the slice level. No volumetric post-processing or 3D reconstruction was applied.

MedSAM was trained on the same axial slices using bounding box prompts derived from the corresponding 2D airway annotations. As with AeroPath, training was initialized from pretrained weights and optimized for 30 epochs, with intermediate checkpoints stored for recovery. Predictions were evaluated directly on 2D slices, consistent with the training setup.

### 2.7 Evaluation protocol

We report Dice and IoU [10]. For paired comparisons, we use the Wilcoxon signed-rank test [34]. We emphasize that overlap metrics alone may not capture clinically relevant properties such as continuity, centerline stability, and boundary distances; we therefore complement evaluation with qualitative overlays [10, 26].

An abstract overview of the complete trachea segmentation pipeline, including data handling, model components, and the hybrid inference strategy, is illustrated in Figure 1.

**Figure 1.**
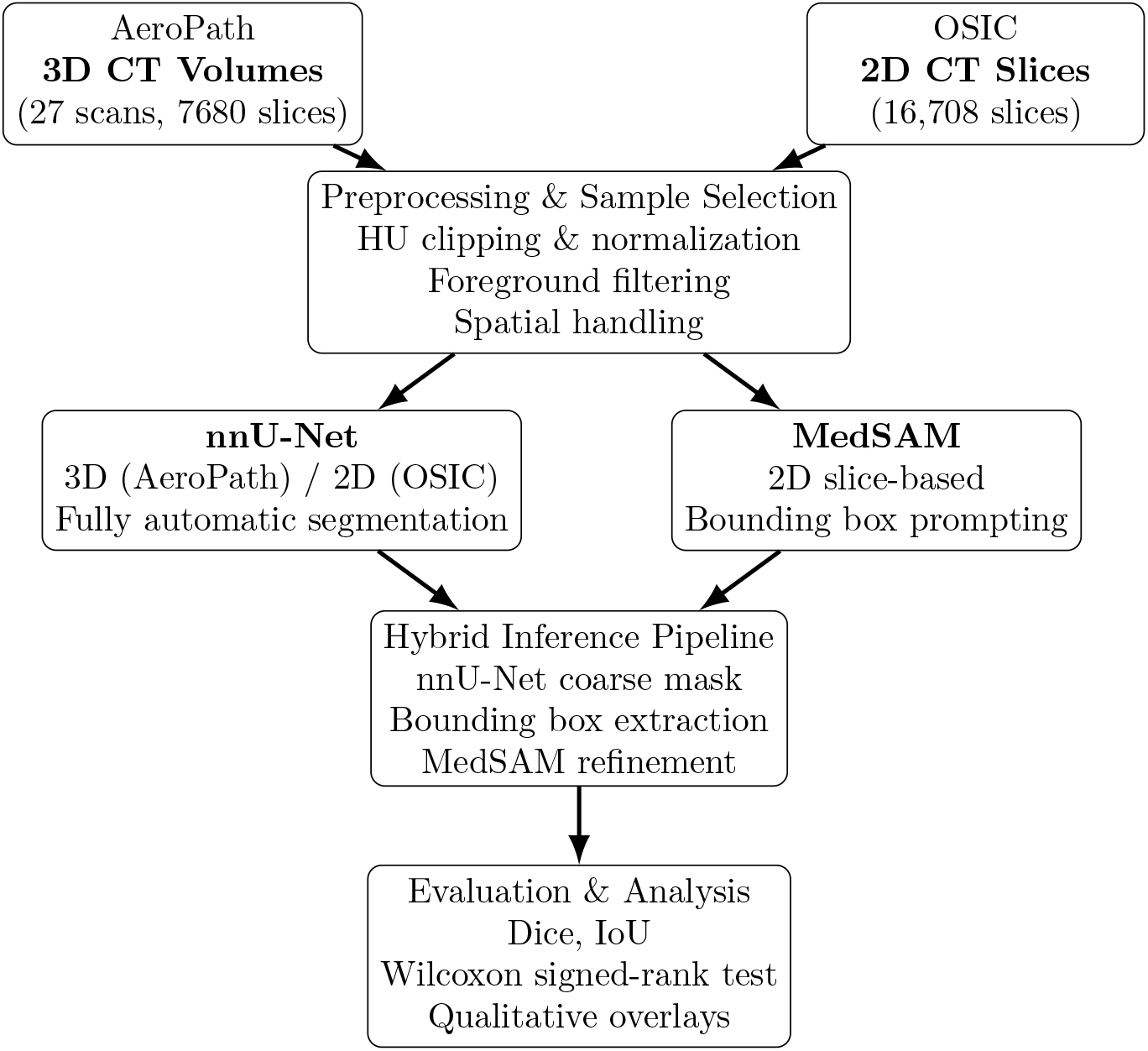
Abstract system overview of the trachea segmentation methodology. Two heterogeneous CT datasets are processed through a shared preprocessing pipeline and segmented using both a fully automatic framework (nnU-Net) and a prompt-conditioned foundation model (MedSAM). A hybrid inference strategy enables fully automatic prompt-based refinement by deriving bounding boxes from nnU-Net predictions. Evaluation combines overlap-based metrics with statistical testing and qualitative assessment.

## 3 Results and Discussion

This section presents quantitative and qualitative results for trachea segmentation on the AeroPath and OSIC datasets and discusses their implications in light of dataset structure, model design, and downstream clinical applicability. Rather than treating results and discussion as disjoint components, empirical findings are interpreted in context to highlight how performance metrics, visual behavior, and deployment considerations interact in practice.

### 3.1 Quantitative Performance Across Datasets

Quantitative segmentation performance for both datasets is summarized in Table 1. Overall, nnU-Net achieves higher overlap scores than MedSAM across both AeroPath and OSIC, with the largest margin observed on AeroPath. This behavior is consistent with the ability of nnU-Net to exploit volumetric continuity and learn spatial context implicitly when consistent inter-slice alignment is available [18, 17]. In volumetric data, elongated anatomical structures such as the trachea benefit substantially from contextual cues across adjacent slices, particularly in anatomically challenging regions where local appearance alone may be ambiguous.

**Table 1:**
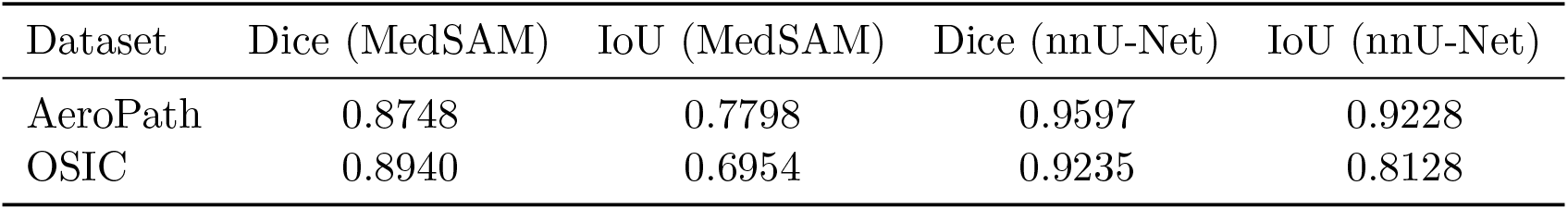
Segmentation performance (Dice and IoU) of MedSAM and nnU-Net on AeroPath and OSIC.

| Dataset | Dice (MedSAM) | IoU (MedSAM) | Dice (nnU-Net) | IoU (nnU-Net) |
| --- | --- | --- | --- | --- |
| AeroPath | 0.8748 | 0.7798 | 0.9597 | 0.9228 |
| OSIC | 0.8940 | 0.6954 | 0.9235 | 0.8128 |

On AeroPath, nnU-Net achieves a Dice score of 0.9597 and an IoU of 0.9228, reflecting both high overlap and strong boundary agreement. MedSAM also performs competitively on this dataset (Dice 0.8748, IoU 0.7798), indicating that prompt-conditioned segmentation can capture the tracheal region reliably when spatial context is stable and prompts are accurate. However, the lower IoU relative to Dice suggests that MedSAM predictions exhibit larger boundary deviations, which disproportionately affect IoU for narrow structures [10].

On OSIC, which lacks reliable volumetric continuity, the performance gap between nnU-Net and MedSAM narrows in terms of Dice (0.9235 vs. 0.8940). This indicates that prompt-based ROI-constrained segmentation remains competitive in strictly slice-based settings. Nevertheless, MedSAM again shows a more pronounced drop in IoU (0.6954), highlighting increased sensitivity to small boundary inconsistencies when operating without volumetric context. These results underscore that Dice alone may overestimate segmentation quality for thin tubular anatomy, while IoU provides complementary insight into boundary fidelity.

Taken together, these quantitative results confirm that self-configuring, fully automatic pipelines provide a strong baseline across heterogeneous regimes [18]. At the same time, they indicate that ROI-constrained prompt-based segmentation remains a viable alternative in settings where volumetric continuity is unavailable or where interpretability and explicit spatial conditioning are prioritized [24].

### 3.2 Qualitative Analysis and Figure-Based Interpretation

Figure 2 presents a representative qualitative example of the segmented trachea in coronal, sagittal, and axial views. The multi-planar visualization highlights the elongated geometry of the trachea, its small axial cross-section, and its close proximity to surrounding mediastinal structures, all of which contribute to segmentation difficulty.

**Figure 2.**
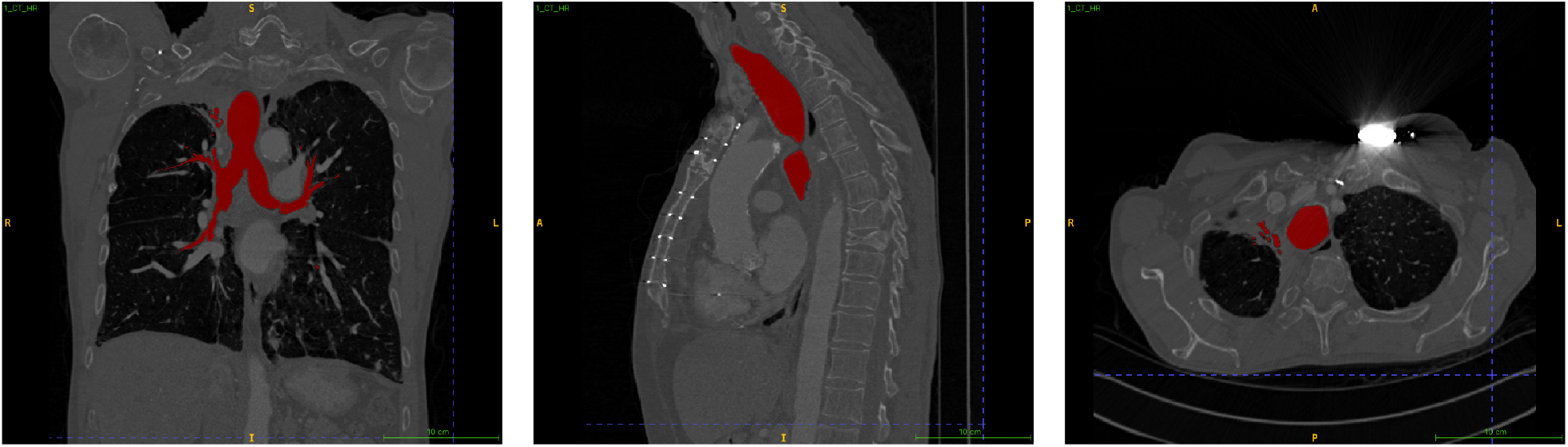
Qualitative example of trachea and proximal airway segmentation shown in three orthogonal CT views. **Left:** coronal view illustrating the elongated cranio–caudal extent of the trachea and its bifurcation into the main bronchi. **Center:** sagittal view highlighting anterior–posterior curvature and spatial relationship to the vertebral column and mediastinal structures. **Right:** axial view demonstrating the small cross-sectional area of the tracheal lumen and its proximity to surrounding tissues. The segmented airway is overlaid in red on the CT images. Orientation markers (R/L/A/P/S/I) follow radiological convention. These views emphasize the anatomical challenges of trachea segmentation, including elongation, small target size, and sensitivity to partial volume effects, which motivate the use of context-aware and ROI-constrained segmentation strategies.

Quantitative metrics alone do not fully capture the behavior of segmentation models on anatomically constrained structures. Figures 3–5 provide qualitative insights that help contextualize the numerical results.

**Figure 3.**
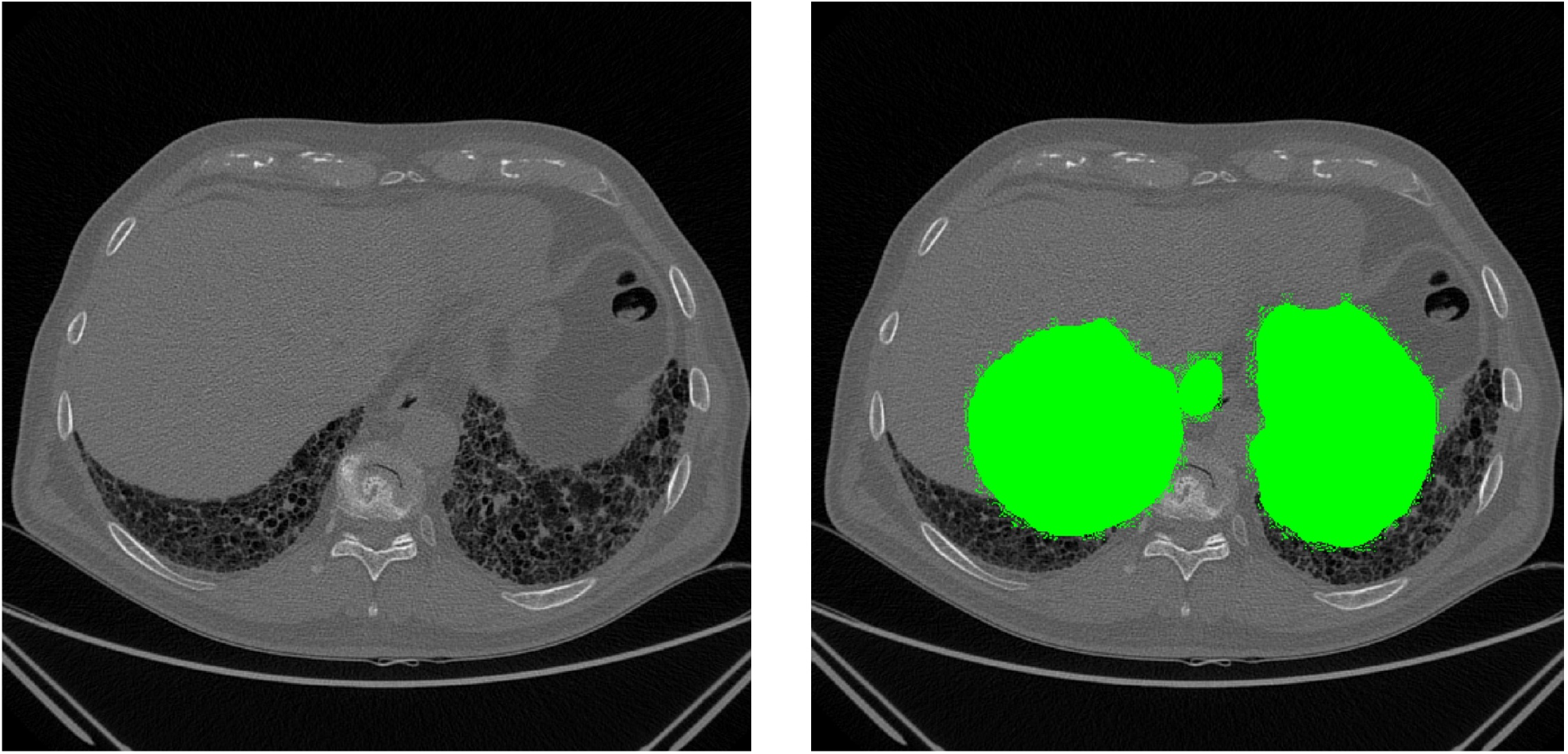
OSIC qualitative example: **raw CT** (left) and **ground-truth overlay** (right). Slice-based datasets lack reliable volumetric continuity, increasing ambiguity from partial volume effects and local contrast variation [29, 7].

**Figure 4.**
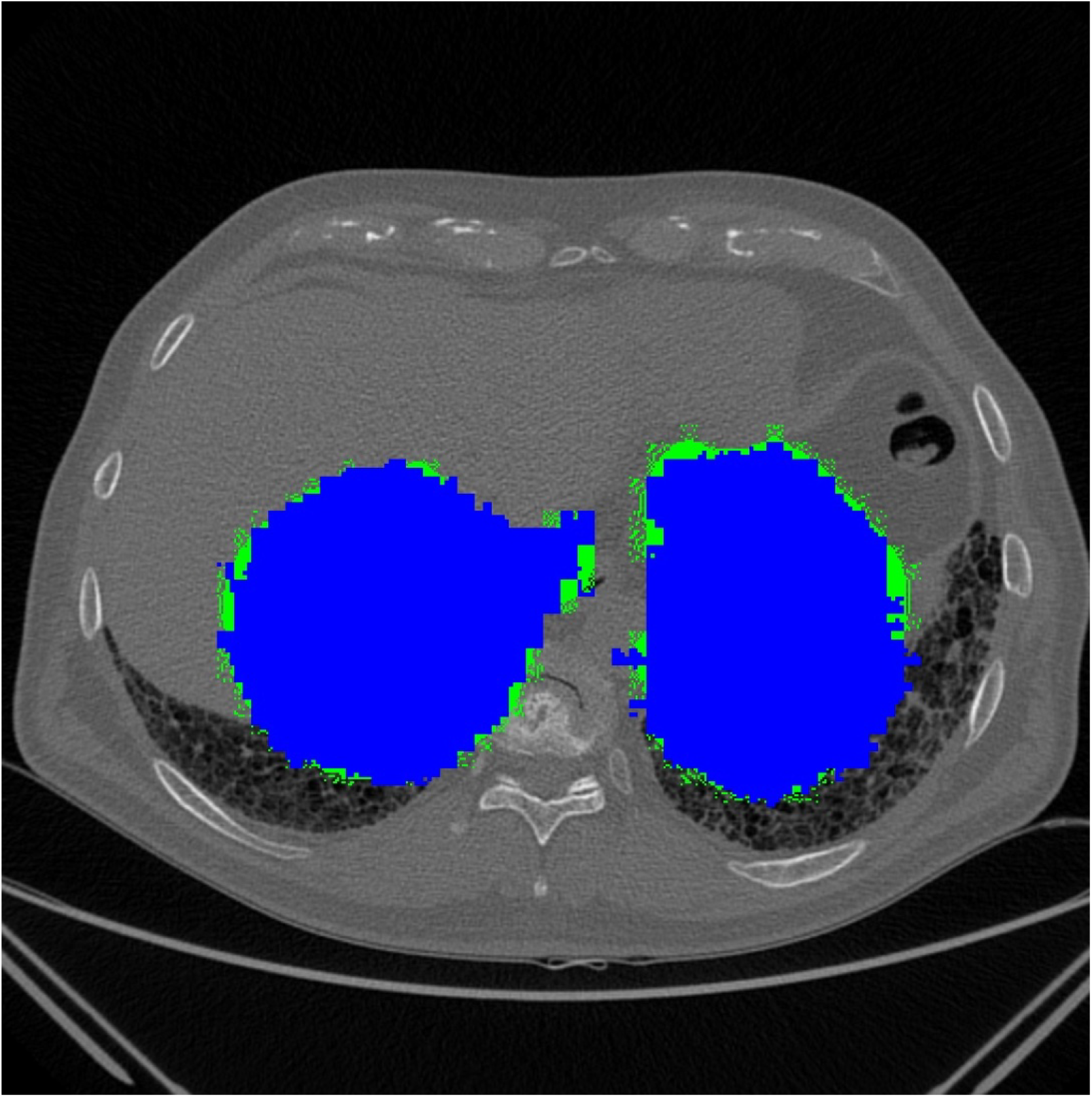
Representative **MedSAM** prediction on OSIC. Prompt-conditioned inference constrains attention to a region-of-interest and can reduce spurious predictions outside the target region, but performance depends on prompt quality and tightness [23, 24].

**Figure 5.**
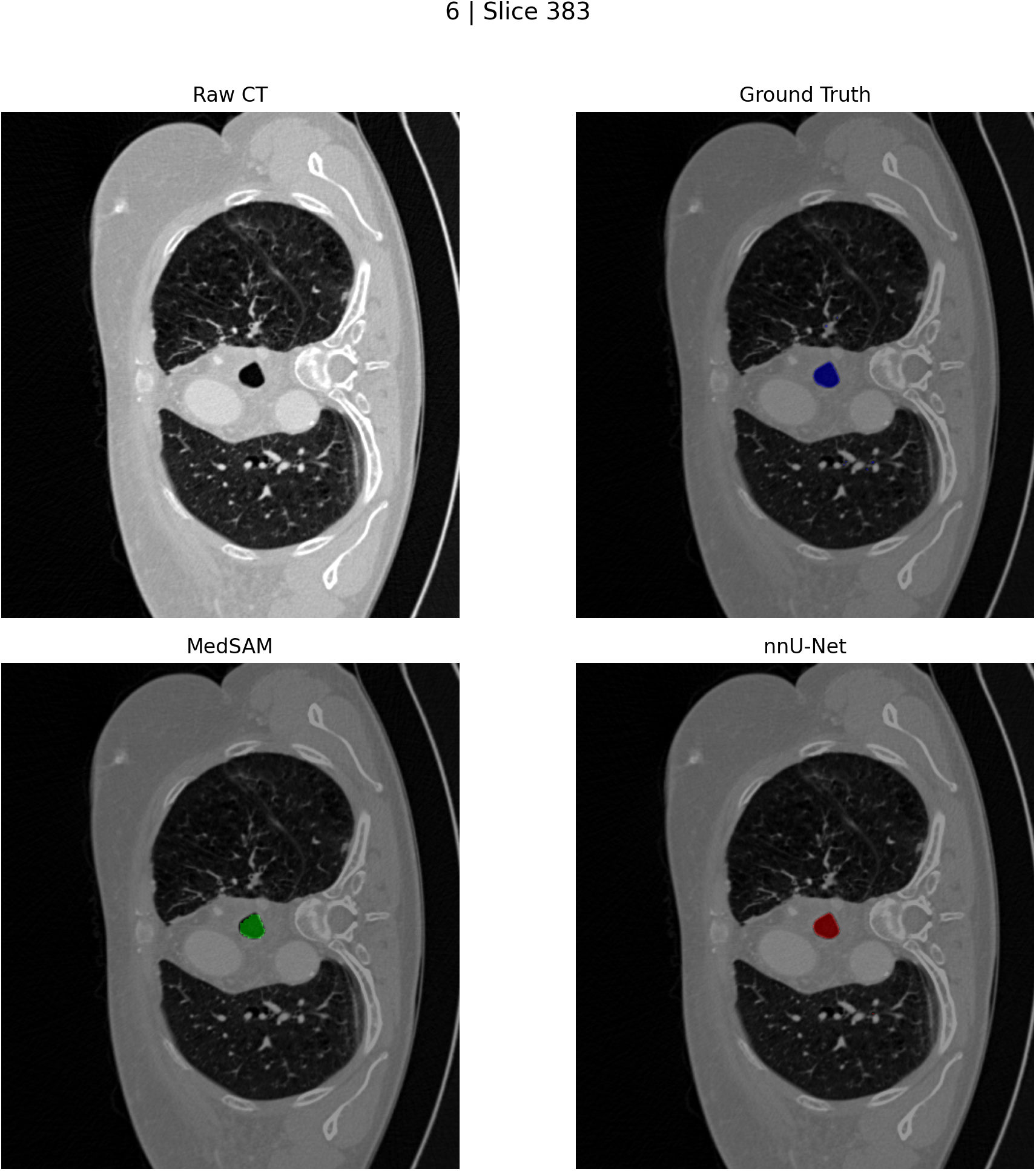
Qualitative comparison (Slice 383). Top: raw CT and ground truth. Bottom: MedSAM and nnU-Net predictions. Differences in boundary adherence can influence downstream airway descriptors (e.g., diameter and cross-sectional area) used in precision airway planning [7].

Figure 3 illustrates a representative OSIC slice, showing the raw CT image alongside the ground-truth trachea annotation. This visualization highlights the inherent difficulty of slice-based trachea segmentation: the target structure occupies a very small fraction of the image, and its appearance may be confounded by adjacent air-filled or low-density regions. Without volumetric continuity, local cues such as lumen shape and intensity gradients must be interpreted in isolation, amplifying ambiguity from partial volume effects and local contrast variation [7, 8].

Figure 4 shows a representative MedSAM prediction on OSIC. The prompt-conditioned formulation constrains attention to a region-of-interest, which helps suppress spurious predictions in irrelevant regions of the image. This behavior is particularly advantageous in slice-based settings, where global context is unreliable. However, the figure also illustrates that segmentation quality remains dependent on prompt tightness: overly permissive prompts can admit adjacent structures, while overly restrictive prompts risk truncating the trachea [23, 24].

Figure 5 provides a direct comparison between ground truth, MedSAM, and nnU-Net predictions on a representative slice. While both models achieve visually plausible segmentations, subtle differences in boundary adherence are evident. These differences may have limited impact on Dice scores but can significantly influence downstream airway descriptors such as diameter profiles, cross-sectional area, and shape regularity—quantities that are critical for precision airway planning [7].

### 3.3 Impact of Dataset Structure on Model Behavior

Across both quantitative and qualitative analyses, dataset structure emerges as a decisive factor influencing segmentation performance, interpretability, and deployment feasibility [9, 26]. In volumetric AeroPath data, spatial continuity enables context-aware learning, favoring fully automatic pipelines that implicitly model anatomical continuity along the tracheal axis. In slice-based OSIC data, the absence of reliable inter-slice context constrains model design and increases sensitivity to local ambiguity, motivating approaches that explicitly incorporate spatial priors.

### 3.4 Fully Automatic Versus Prompt-Based Segmentation

Fully automatic segmentation with nnU-Net demonstrates strong robustness and scalability across both datasets, reflecting its ability to jointly learn localization and segmentation without auxiliary inputs [18, 11]. This property is particularly valuable in high-throughput clinical settings where manual interaction or prompt specification is infeasible.

Prompt-based segmentation with MedSAM offers a complementary paradigm. By conditioning segmentation on explicit spatial cues, MedSAM enables anatomically constrained predictions and improved interpretability, especially in slice-based regimes [24, 23]. However, this flexibility comes at the cost of prompt dependency, introducing an additional source of variability that must be managed carefully.

### 3.5 Hybrid Prompting for Practical Deployment

The hybrid inference strategy reconciles these trade-offs by combining nnU-Net for automatic localization with MedSAM for ROI-constrained refinement. This design preserves full automation while leveraging the strengths of foundation models for anatomically focused segmentation. Beyond performance considerations, the explicit prompt also provides a transparent “focus region” that can aid clinician-facing review, debugging, and trust calibration [35, 36].

### 3.6 Quantitative Metrics Versus Clinical Plausibility

Finally, the results highlight the limitation of relying solely on overlap-based metrics. Dice and IoU do not fully capture clinically relevant properties such as boundary distance, centerline stability, and continuity, which are essential for precision airway management [10]. The qualitative figures demonstrate that models with similar overlap scores may differ meaningfully in local geometry, reinforcing the need for evaluation protocols aligned with downstream clinical use.

### 3.7 Implications for Precision Airway Workflows

From a deployment perspective, the findings suggest that dataset characteristics should guide model selection. Consistent volumetric CT favors context-aware fully automatic pipelines, whereas heterogeneous or slice-based archives may benefit from ROI-constrained prompt-based refinement when reliable automatic prompting is available. These observations align with broader recommendations that clinical AI systems balance performance, robustness, and interpretability to support safe and effective deployment [37, 38].

## 4 Limitations

Despite demonstrating the feasibility and robustness of both fully automatic and prompt-based approaches for trachea segmentation, this study has several limitations that merit careful consideration.

A primary limitation arises from the heterogeneous nature of the datasets used for evaluation. The analysis was restricted to two datasets with fundamentally different structural properties: a volumetric dataset with consistent spatial alignment and a large-scale slice-based dataset lacking reliable inter-slice continuity. While this heterogeneity increases ecological validity and reflects realistic clinical data acquisition regimes, it inevitably constrains the ability to conduct tightly controlled comparisons under identical data representations. Differences in spatial organization, preprocessing requirements, and indexing conventions can also complicate visualization, traceability of predictions to original image coordinates, and reproducibility of qualitative assessment across datasets [25, 26]. Consequently, some observed performance differences may reflect dataset-specific constraints rather than intrinsic model capabilities.

A second limitation concerns the sensitivity of prompt-conditioned segmentation to prompt quality. MedSAM relies on explicit spatial prompts at inference time, and although bounding box prompts provide a comparatively robust and interpretable spatial prior for elongated anatomical structures such as the trachea, they remain susceptible to error. In the proposed hybrid pipeline, bounding boxes are derived automatically from coarse nnU-Net predictions, which introduces the possibility of error propagation: inaccuracies in the initial localization may be transferred directly to the refinement stage [24]. Overly loose prompts may permit the inclusion of adjacent air-filled or low-density structures, while overly tight prompts may truncate the trachea, particularly in slices affected by partial volume effects or weak contrast. This dependency is inherent to prompt-based segmentation paradigms and highlights the importance of robust, uncertainty-aware prompt generation strategies.

Third, the evaluation protocol focused primarily on overlap-based metrics, namely Dice similarity coefficient and Intersection-over-Union. Although these measures are widely adopted and provide a useful first-order assessment of segmentation quality, they do not fully capture the clinical requirements of trachea segmentation for airway assessment and planning. Clinically relevant properties such as boundary accuracy, centerline deviation, geometric continuity, topological correctness, and sensitivity to localized boundary errors—particularly near anatomically critical regions such as the tracheal inlet and carina—are not adequately reflected by global overlap scores [10]. As a result, models with similar Dice values may differ substantially in their suitability for downstream clinical use, especially when derived measurements such as diameter profiles or cross-sectional areas are considered.

Finally, the target clinical cohort intended for downstream deployment and integration with decision-support workflows has not yet been fully consolidated. The experiments presented here were designed to evaluate methodological behavior and robustness under heterogeneous and realistic conditions rather than to optimize performance for a single institution-specific data distribution. Consequently, the reported results should be interpreted as methodological validation rather than as final deployment-ready performance. Dataset-aware refinement, recalibration, and external validation will be required once the final clinical cohort is defined, in order to minimize distribution shift and ensure reliable generalization in the intended clinical setting.

## 5 Future Work

Future work will be directed toward improving robustness, evaluation relevance, and deployment readiness of trachea segmentation within precision airway management workflows. Model refinement will be performed once the finalized clinical cohort is consolidated. In particular, nnU-Net will be reconfigured to match the final voxel spacing, intensity distribution, and annotation conventions of the target dataset, leveraging its self-configuring design. In addition to the standard nnU-Net framework, derivative variants (e.g., nnU-Netv2 and lightweight adaptations optimized for inference efficiency) will be explored to assess trade-offs between accuracy, computational cost, and deployment feasibility.[18]

For prompt-based segmentation, MedSAM will be fine-tuned according to cohort-specific prompt distributions and labeling practices. Beyond the baseline MedSAM configuration, recent and emerging derivatives of SAM adapted for medical imaging—such as domain-adaptive SAM variants, parameter-efficient fine-tuning approaches, and lightweight prompt-conditioned decoders—will be investigated to improve specialization for thin, elongated anatomical structures like the trachea while maintaining generalization.[24]

To reduce sensitivity to prompt quality, uncertainty-aware and multi-prompt strategies will be investigated. These include probabilistic bounding boxes, sequential prompting schemes, and prompt ensemble techniques. Such approaches are expected to mitigate failure modes arising from incomplete or noisy coarse segmentations and to support robustness calibration under heterogeneous imaging conditions.[39, 40]

Evaluation will be extended beyond overlap-based metrics to include clinically grounded measures. Performance will be assessed using boundary-distance metrics (e.g., Average Surface Distance and Hausdorff distance), centerline-based deviations, continuity and topology checks, and region-specific analyses (e.g., upper trachea versus carina).[10] Importantly, it will be validated whether improvements in segmentation quality translate into stable and reliable downstream airway descriptors—such as diameter profiles, lumen consistency, and geometric proxies—that are directly relevant to clinical planning.

Finally, segmentation-derived anatomical descriptors will be integrated with structured Electronic Health Record (EHR) variables to enable downstream prediction and decision-support tasks for precision airway management. This multimodal integration is expected to support patient-specific risk stratification, procedural planning, and longitudinal assessment within AI-assisted clinical workflows.[41, 36]

## 6 Conclusion

We presented a systematic study of trachea segmentation from CT imaging using two complementary paradigms: a fully automatic segmentation framework (nnU-Net) and a prompt-conditioned foundation model (MedSAM). By evaluating these approaches on both volumetric (AeroPath) and slice-based (OSIC) datasets, we showed that dataset structure strongly influences segmentation reliability, interpretability, and deployment feasibility. Fully automatic nnU-Net provides robust and scalable performance, particularly when volumetric continuity enables exploitation of spatial context. Prompt-based MedSAM offers anatomically constrained segmentation through explicit spatial priors and facilitates interpretability, but introduces sensitivity to prompt quality. A hybrid inference strategy—using nnU-Net for automatic localization and MedSAM for prompt-conditioned refinement—offers a practical pathway to integrate foundation models into fully automatic pipelines without sacrificing scalability. Overall, our findings suggest that foundation models can complement classical medical segmentation methods, especially for anatomically constrained tasks, when paired with reliable automatic prompting and evaluated with metrics aligned to clinical utility. Future work will refine the pipeline on the finalized clinical cohort and integrate segmentation-derived descriptors with EHR data to support downstream precision airway assessment and decision support.

## Acknowledgments

This work was funded by the European Union Recovery and Resilience Facility of the NextGenerationEU instrument, through the Research and Innovation Foundation (Project: Improving Personalized Medicine via AI-based Precision Tracheostomy-PRECIOUS, ENTERPRISES/0223/Sub-Call1/0263)

## Notes

### Competing Interest Statement

The authors have declared no competing interest.

### Summary of Updates

In this revision, we have made limited but important updates to the manuscript. Specifically: Founding Information: We have clarified and updated the founding details to ensure accuracy and completeness. Author List Update: The author list has been revised to reflect correct contributions and affiliations. No other changes have been made to the scientific content, results, or conclusions of the manuscript.

## References

[1] MW El-Anwar, AA Nofal, MA Shawadfy, A Maaty, and Khazbak AO. Tracheostomy in the intensive care unit: indications, timing, technique, and outcomes. International archives of otorhinolaryngology, 21(1):33–37, 2017.

[2] Christian Putensen, Niklas Theuerkauf, Ulrich Guenther, Mario Vargas, and Paolo Pelosi. Percutaneous tracheostomy in the icu: a review of indications, techniques and complications. Critical Care, 18:544, 2014.

[3] Donald E. G. Griesdale, T. L. Bosma, T. Kurth, G. Isac, and D. R. Chittock. Complications of endotracheal intubation in the critically ill. Intensive Care Medicine, 34:1835–1842, 2008.

[4] DT Wong and A. Kumar. Endotracheal tube malposition in adults: clinical consequences and prevention. Canadian journal of anaesthesia = Journal canadien d’anesthesie, 2006.

[5] Rüdiger R Noppens. Airway management in the icu. Acta Clin Croat., 2012.

[6] PC Pacheco-Lopez, LC Berkow, AT Hillel, and LM. Akst. Complications of airway management: a review. Respir Care, 2014.

[7] Guillaume Chassagnon, Maria Vakalopoulou, Nikos Paragios, and Marie-Pierre Revel. Artificial intelligence applications for thoracic imaging. European Journal of Radiology, 129:109118, 2020.

[8] Pechin Lo, Jon Sporring, Haseem Ashraf, Jesper J. H. Pedersen, and Marleen de Bruijne. Vessel-guided airway tree segmentation: A voxel classification approach. Medical Image Analysis, 16(4):807–818, 2012.

[9] Tobias Heimann and Hans-Peter Meinzer. Statistical shape models for 3d medical image segmentation: a review. Medical Image Analysis, 13(4):543–563, 2009.

[10] Abdel Aziz Taha and Allan Hanbury. Metrics for evaluating 3d medical image segmentation: analysis, selection, and tool. BMC Medical Imaging, 15:29, 2015.

[11] Geert Litjens, Thijs Kooi, Babak E. Bejnordi, Arnaud A. A. Setio, Francesco Ciompi, Mohsen Ghafoorian, Jeroen van der Laak, Bram van Ginneken, and Clara I. Sánchez. A survey on deep learning in medical image analysis. Medical Image Analysis, 42:60–88, 2017.

[12] Dinggang Shen, Guorong Wu, and Heung-Il Suk. Deep learning in medical image analysis. Annual Review of Biomedical Engineering, 19:221–248, 2017.

[13] Jonathan Long, Evan Shelhamer, and Trevor Darrell. Fully convolutional networks for semantic segmentation. CVPR, 2015.

[14] Olaf Ronneberger, Philipp Fischer, and Thomas Brox. U-net: Convolutional networks for biomedical image segmentation. In xMICCAI, 2015.

[15] Zongwei Zhou, Mohammad R. Siddiquee, Nima Tajbakhsh, and Jianming Liang. Unet++: A nested u-net architecture for medical image segmentation. In DLMIA, 2018.

[16] Ozan Oktay, Jo Schlemper, et al. Attention u-net: Learning where to look for the pancreas, 2018.

[17] Fausto Milletari, Nassir Navab, and Seyed-Ahmad Ahmadi. V-net: Fully convolutional neural networks for volumetric medical image segmentation. In 3DV, 2016.

[18] Fabian Isensee, Paul F. Jaeger, Simon A. A. Kohl, Jens Petersen, and Klaus H. Maier-Hein. nnu-net: a self-configuring method for deep learning-based biomedical image segmentation. Nature Methods, 18(2):203–211, 2021.

[19] Jieneng Chen, Yufan Lu, Qihang Yu, et al. Transunet: Transformers make strong encoders for medical image segmentation, 2021.

[20] Ali Hatamizadeh, Yi Tang, et al. Swin unetr: Swin transformers for semantic segmentation of brain tumors in mri images. In MICCAI, 2022.

[21] Ze Liu, Yutong Lin, Yue Cao, et al. Swin transformer: Hierarchical vision transformer using shifted windows. In ICCV, 2021.

[22] Liang-Chieh Chen, Yukun Zhu, George Papandreou, Florian Schroff, and Hartwig Adam. Encoder-decoder with atrous separable convolution for semantic image segmentation. In ECCV, 2018.

[23] Alexander Kirillov, Eric Mintun, Nikhila Ravi, Hanzi Mao, Chloe Rolland, Laura Gustafson, Tete Xiao, Spencer Whitehead, Alexander C. Berg, Piotr Dollár, and Ross Girshick. Segment anything, 2023.

[24] Jun Ma, Bo Wang, et al. Segment anything in medical images, 2023.

[25] Lena Maier-Hein, Martin Eisenmann, Annika Reinke, et al. Why rankings of biomedical image analysis competitions should be interpreted with care. Nature Communications, 9:5217, 2018.

[26] Annika Reinke et al. Common pitfalls and recommendations for evaluating and reporting on medical image analysis. Medical Image Analysis, 73:102230, 2021.

[27] A. Panayides, H. Chen, N. Filipovic, T. Geroski, J. Hou, K. Lekadir, K. Marias, G. Matsopoulos, Giorgos Papanastasiou, P. Sarder, G. Tourassi, S. Tsaftaris, H. Fu, E. Kyriacou, Christos Loizou, M. Zervakis, J.H. Saltz, F. Shamout, K. Wong, and M.S. Pattichis. Position paper: Artificial intelligence in medical image analysis: Advances, clinical translation, and emerging frontiers. IEEE Journal of Biomedical and Health Informatics, PP:1–16, 12 2025.

[28] Nikolaos Papandrianos et al. Aeropath: Benchmark dataset for airway segmentation and analysis, 2024.

[29] Open Source Imaging Consortium (OSIC). Osic pulmonary fibrosis progression (kaggle competition dataset). Kaggle, 2020.

[30] Mateusz Buda, Ashirbani Saha, and Maciej A. Mazurowski. Deep learning in radiology: An overview of the concepts and a survey of the state of the art with focus on mri. Medical Image Analysis, 73:102145, 2021.

[31] Carole H. Sudre, Wenqi Li, Tom Vercauteren, Sebastien Ourselin, and M. Jorge Cardoso. Generalised dice overlap as a deep learning loss function for highly unbalanced segmentations. In DLMIA, 2017.

[32] Diederik P. Kingma and Jimmy Ba. Adam: A method for stochastic optimization, 2015.

[33] Ian Goodfellow, Yoshua Bengio, and Aaron Courville. Deep learning, 2016.

[34] Frank Wilcoxon. Individual comparisons by ranking methods. Biometrics Bulletin, 1(6):80–83, 1945.

[35] Finale Doshi-Velez and Been Kim. Towards a rigorous science of interpretable machine learning, 2017.

[36] Eric Topol. High-performance medicine: the convergence of human and artificial intelligence. Nature Medicine, 25:44–56, 2019.

[37] Christopher J. Kelly, Alan Karthikesalingam, Mustafa Suleyman, Greg Corrado, and Dominic King. Key challenges for delivering clinical impact with artificial intelligence. BMC Medicine, 17:195, 2019.

[38] Andre Esteva, Alexandre Robicquet, Bharath Ramsundar, Volodymyr Kuleshov, Mark DePristo, Katherine Chou, Claire Cui, Greg Corrado, Sebastian Thrun, and Jeff Dean. A guide to deep learning in healthcare. Nature Medicine, 25:24–29, 2019.

[39] Xiangbin Liu, Liping Song, Shuai Liu, and Yudong Zhang. A review of robustness in deep learning for medical image segmentation. Sustainability, 2021.

[40] Ian J. Goodfellow, Jonathon Shlens, and Christian Szegedy. Explaining and harnessing adversarial examples, 2015.

[41] Alvin Rajkomar, Eyal Oren, Kai Chen, et al. Scalable and accurate deep learning with electronic health records. npj Digital Medicine, 1:18, 2018.

